# High-throughput Microfluidic 3D Outer Blood-Retinal Barrier Model in a 96-Well Format: Analysis of Cellular Interactions and Barrier Function in Retinal Health and Disease

**DOI:** 10.1101/2023.12.01.569537

**Authors:** Jiho Kim, Youngsook Song, Amber L. Jolly, Taeseon Hwang, Suryong Kim, Byungjun Lee, Jinhwan Jang, Dong Hyun Jo, Kyusuk Baek, Tsung-Li Liu, Sanghee Yoo, Noo Li Jeon

## Abstract

Numerous diseases, including AMD (age-related macular degeneration), arise from the blood-retinal barrier and blood vessel abnormalities in the eye; unfortunately, there is a lack of reliable in-vitro models for their systematic study. This study describes a high-throughput microphysiological system (MPS) designed to model the outer Blood-Retinal Barrier (oBRB). The MPS platform is engineered to integrate seamlessly with high-content screening technologies, utilizing a design with a single oBRB model incorporating RPE (retina pigment epithelial cells) and endothelial cell co-culture to fit within a single 96-well. Arranged in the standard 96-well plate format, the platform allows high-throughput assessment of barrier integrity through 3D confocal imaging (ZO-1 staining), Trans Epithelial Electrical Resistance (TEER), and permeability measurements.

The oBRB model enables the investigation of crosstalk among different cell types in co-culture. This includes assessing changes in the barrier integrity of the Retinal Pigment Epithelium (RPE) monolayer and investigating neovascularization events resulting from endothelial cell remodeling. The platform is positioned for utility in drug discovery and development efforts targeting diseases involving oBRB damage and choroidal neovascularization, such as age-related macular degeneration (AMD).

## 1. Introduction

Age-related Macular Degeneration (AMD) is a prominent cause of irreversible vision loss among elderly populations globally[1]. The dysfunction and subsequent breach of the blood-retinal barrier (BRB) accelerates the influx of inflammatory and angiogenic factors into the retina[2], thereby contributing to neovascularization and degenerative processes characteristic of wet and dry AMD, respectively. In wet AMD, the BRB plays a pivotal role in maintaining retinal homeostasis and is crucially implicated in AMD’s pathogenesis[3]. During the pathogenesis of AMD, it is first observed that the outer Blood-Retinal Barrier (oBRB) becomes damaged due to cumulative hypoxic, oxidant, or inflammatory stress[4–6]. Subsequently, in the vicinity of the Retinal Pigment Epithelium (RPE), there is an increase in the secretion of Vascular Endothelial Growth Factor (VEGF) and a decrease in Pigment Epithelium-Derived Factor (PEDF)[7]. In this process, the damaged RPE induces new vessel formation in the choroid and makes these induced vessels leaky, thereby accelerating the dysfunction of the RPE[8].

3D Microphysiological systems (MPS) are well suited for exploring the mechanical intricacies of diseases like AMD in a controlled and replicable laboratory setting. The value of MPS in modeling BRB dysfunction lies in its potential to mimic the pathophysiological conditions in vitro, providing insights into the cellular and molecular mechanisms underpinning the disease and concurrently serving as a platform for pharmacological testing. Recent studies have leveraged MPS to model AMD, yet these models fall short in accurately mimicking blood-retinal barrier (BRB) pathology and in evaluating retinal pigment epithelium (RPE) barrier functions. Most of these MPS oBRB models examined co-cultures of endothelial cells and RPEs cultured in proximity and often did not include results of AMD-inducing treatments as part of the study[9–11]. The most advanced in vitro model of the outer blood-retinal barrier to date was published using 3D Bioprinting, which included iPSC-RPEs on the top and a capillary bed of iPSC-endothelial cells, pericytes, and fibroblasts on the bottom of a biodegradable PLGA membrane[12]. This approach allowed the measurement of crosstalk of multiple cell types, including the influence of RPE cells on fibroblasts to induce fibroblasts to secrete components critical to forming an in vivo-like Bruch’s membrane. Unfortunately, this complicated system is not useful for HTS and, for this reason, is not readily amenable to drug screening and development efforts. Limitations in throughput and an absence of drug response data compromise their utility for clinical prediction and drug screening[13–15].

In this research, we unveil a microphysiological system adeptly recapitulating normal oBRB and its degradation to presenting BRB pathology facilitated by a damage inducer. The developed model uniquely allows for the quantifiable evaluation of the barrier function via three distinct methodologies: Trans Epithelial Electrical Resistance (TEER), tight junction protein ZO-1 staining, and a permeability assay of the RPE layer. Moreover, our system was employed to test efficacious drugs pertinent to block neovascularization, exemplified by Bevacizumab, an anti-VEGF drug currently foregrounding wet AMD therapeutic strategy. Through meticulous experimentation, our model not only endorsed the potential for a nuanced understanding of BRB dysregulation and pathological processes but also established itself as a potent tool for future drug screening and development endeavors, underlining a substantial advancement in the realm of ophthalmological research.

## 2. Result

### 2.1 Design of Curio-Barrier Microphysiological System Supporting Barrier Formation

We have developed an MPS organ chip to simulate the epithelial barrier with high reproducibility and reliability. This organ chip was designed in a 96-well plate format, considering compatibility with the SBS format for automated devices supporting High Content Screening (HCS) or High Throughput Screening (HTS) (Fig1.A). For mass production, a polystyrene plate body was manufactured through injection molding. The plate body can be easily attached to a bottom polycarbonate plate using an adhesive film for assembly (Fig1.B). Each well of the plate consists of three open channels and two reservoirs. The middle channel with the lowest height was patterned first using a hydrogel. Following the middle channel patterning, the Side1 channel on the left was patterned second, which is higher than the middle channel. Both channels have a rail structure with a ceiling overhead for precise patterning geometry. The Side2 channel on the right is the last channel for patterning, which has a narrow valley-like structure (Fig1.C). After patterning the three channels, flow within the system can be generated from the two reservoirs through hydrostatic pressure.

**Fig. 1.**
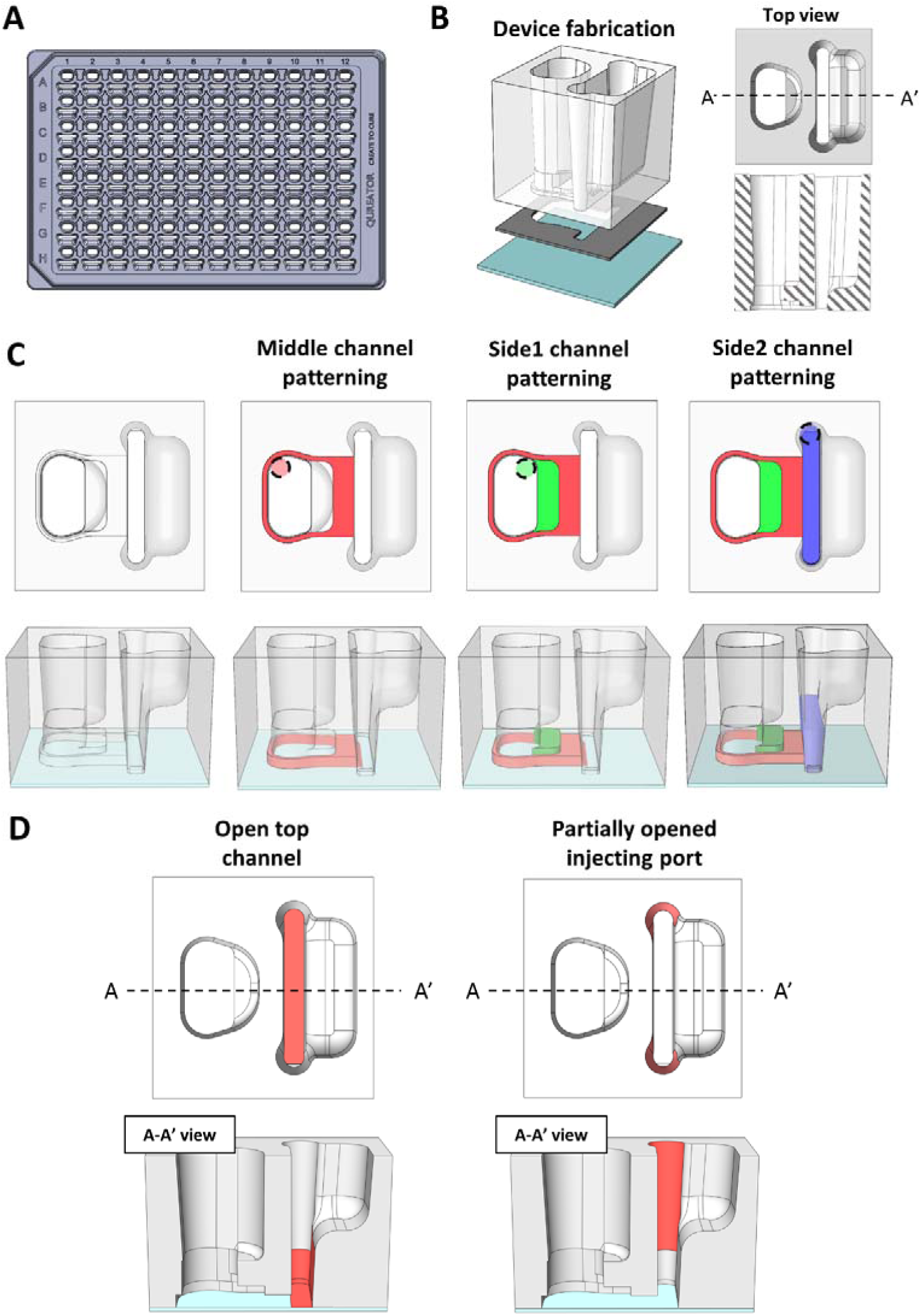
Overview of the device for Curio-Barrier MPS. A) Illustration of the 96well plate used in the experiment. B) Fabrication and the design of the detail well of the device. C) Procedure of the patterning of the device. D) Key features of the device for stable patterning and robust cell culture.

This design incorporates additional features to generate a reliable barrier formation (Fig1.D). 1) The narrow valley-like structure in the Side2 channel allows consistent barrier formation without additional actions, such as tilting the chip. Moreover, the Side2 channel has an open-top design, which facilitates the easy removal of any bubbles that might form during patterning, thereby enhancing the stability of the barrier formation. 2) The partially opened injection port reduces pressure when removing the pipette after fluid patterning, ensuring consistent cell distribution within the channel. 3) The curved structure slows the fluid velocity on a hydrophilic surface, ensuring the fluid is selectively patterned in the desired direction without dispersion.

### 2.2 Engineering the Blood-Retinal Barrier Models in Curio-Barrier MPS

The valley-shaped Side2 channel can create a stable monolayer or multilayer of different cell types over the hydrogel surface. For multilayer formation, the fluid can be removed by suctioning within the Side2 channel, and other cell types can be added in the same channel for the second layer (Fig2.A). Through such repetitive serial patterning, one can culture multilayered tissue structures without needing a Transwell membrane. We established the oBRB using a monolayer and inner BRB using double layers to demonstrate the feasibility of both mono and multilayer culturing in this platform (Fig2.B).

**Fig. 2.**
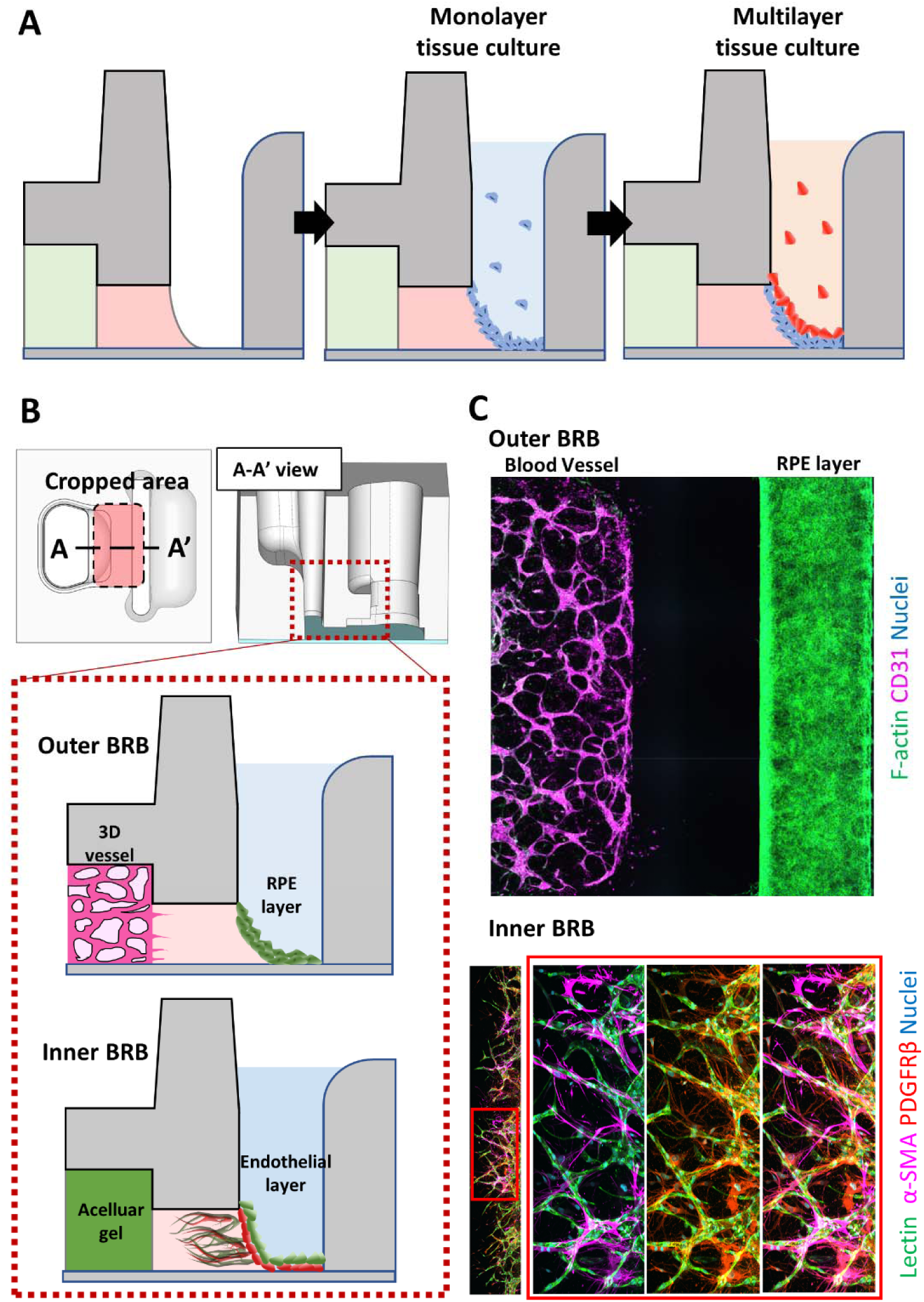
Curio-Barrier supporting epithelial monolayer or multilayer of the blood-retinal barrier model. A) Sequential patterning and culturing the monolayer of cells to multi-layer of different cell types. B) Illustration of the outer BRB of the monolayer and Inner BRB of double layers inside the device. C) Immunostaining results of oBRB and iBRB model. RPE cells (F-actin) and endothelial cells (CD31) cultured in oBRB model. Endothelial cells (Lectin) and pericytes (α-SMA, PDGFRβ) cultured in iBRB model.

For the oBRB, an acellular gel was patterned in the middle channel. In the Side 1 channel on the left, EC (endothelial cells) and the hydrogel mixture were patterned to enable the formation of 3D vessels. Subsequently RPE cells were added to the Side2 channel on the right and the RPE monolayer was formed along the side slope of the hydrogel (Fig2.C). In the case of the inner BRB, it is vital to emulate the configuration of pericytes enveloping vessels [16]. Therefore, we constructed a double-layer configuration; the pericyte layer was added to the slope in the Side 2 channel; after ensuring their stable anchorage in the gel, the endothelial monolayer was serially added where pericytes encased the endothelial tubes during angiogenesis (fig2.C). We confirmed the capability of developing a monolayer of oBRB and implementing multilayer structures of iBRB in Curio-Barrier MPS.

### 2.3 Evaluation of Damage Induction in the Normal oBRB

For the construction of oBRB, an acellular fibrin gel was placed in the middle channel, the EC and fibrin gel mixture was loaded into the Side1 channel, and the Side2 channel was filled with medium. On the second day, the medium was removed, and RPE cell suspension was added, allowing them to settle firmly along the slope of the gel. The pre-wetted channel helped reduce bubble formation between the gel and cell suspension. To implement the normal oBRB emulating the healthy state for both EC and RPE, we used the EGM-2 medium in the EC side reservoir (Side 1) and the RPE differentiation medium in the RPE side reservoir (Side 2). After loading the RPE layer, the culture was continued for an additional 6 days, totaling 7 days.

The barrier function of the oBRB was verified using the three most commonly employed methods that are capable of quantifying whether tight junctions were formed adequately in the RPE monolayer[17] [18]: The permeability assay using FITC-70 kDa dextran in the normal oBRB showed stable tight junction formation with almost no diffusion occurring over 30 minutes (Suppl.Fig1). Additionally, the expression of ZO-1 was confirmed through immunostaining. Its hexagonal morphology between RPE cells was verified (Fig.3B). The resistance value measured using the TEER equipment also significantly increased, yielding a TEER value of approximately 84.1 Ω·cm² compared to the control where the RPE barrier was absent (Fig.3.C).

**Fig. 3.**
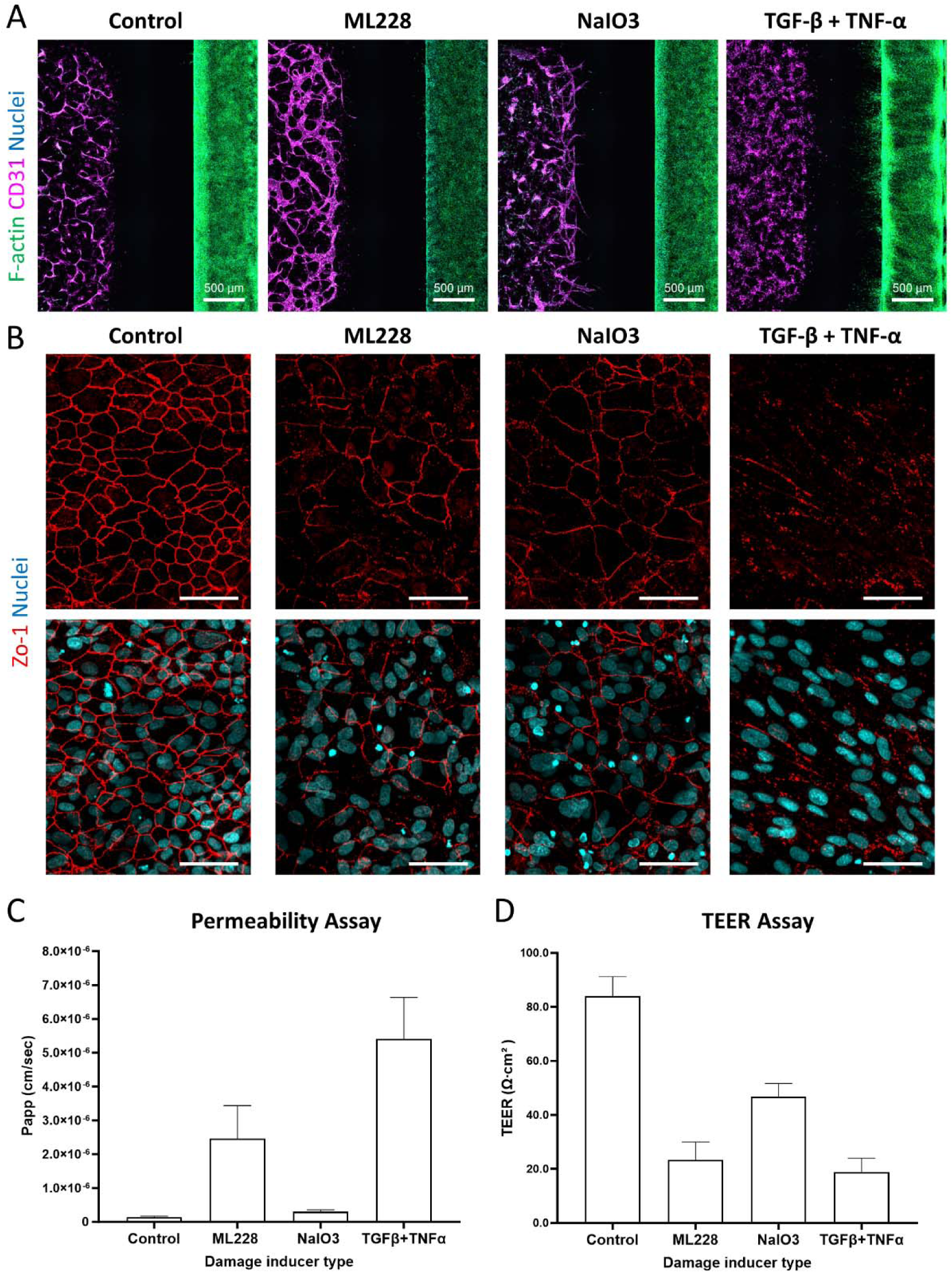
Morphological changes of the RPE layer and vascular network by damaging inducers A) Immunostaining results of RPE cells (F-actin) and endothelial cells (CD31) in the normal or each damage inducer treated oBRB (B) Disappearing Zo-1 intensity by each damage inducer treatment (C) Quantification of 70 kDa FITC-dextran permeability assay showing an increase of permeability by each damage inducer D) TEER value decreasing by each damage inducer treatment

AMD is associated with the deterioration of RPE cells over the course of a lifetime, and a variety of factors can contribute to this damage. Notable inducers of RPE damage include oxidative stress, hypoxic stress, and inflammatory stress [8]. Therefore, we treated the oBRB model with damaging inducers to mimic a stressed RPE state and examined the effects on the integrity and barrier function of the RPE monolayer. To test each stress factor within our platform, hypoxic stress was induced using ML228 [19], oxidative stress using NaIO₃ [20], and a mixture of TNF-α and TGF-β, known as inflammatory stressors, was treated [21, 22]. Each damage inducer was applied for 48 hours after seeding the RPE cells, and the cells were treated twice at 48-hour intervals.

All three damaging inducers decreased the barrier function while distinctly altering the morphology of the blood vessels and RPE cells. ML228 caused a general decrease in F-actin expression in the RPE layer. In contrast, the vessels appeared generally thicker than the vessels in the control and the group treated with NaIO₃ (Fig.3.A). The NaIO₃ treatment decreased the F-actin intensity compared to the control, but higher than the intensity in the ML228-treated group. New blood vessel sprouting was observed in the NaIO₃-treated oBRB model, which appears to be similar to choroidal neovascularization. The TNF-α and TGF-β treatment resulted in the most dramatic damage, as evident from the immunostaining results. The RPE collapsed, failing to form layers and extending towards the middle channel. Notably, different responses to the inducer were observed between the boundary and its center of the RPE channel. Although an overall increase in CD31 was noted in the blood vessels, the vascular network appeared disrupted.

Regarding the immunostaining results of the tight junction protein ZO-1, changes in the expression pattern were observed compared to the healthy hexagonal morphology in the untreated oBRB, corresponding to the morphological alterations in the RPE by the damaging inducers. All groups treated with damage inducers exhibited a marked decrease in intensity and loss of clear hexagonal morphology (Fig3.B). Even in the case of NaIO₃ treatment, where the morphological changes in F-actin and blood vessels were relatively minor, the hexagonal shape started disappearing. In the group treated with TNF-α and TGF-β, which showed severe morphological changes, a complete loss of the hexagonal shape of ZO-1 was evident.

The results of the permeability and TEER measurements were consistent with the morphological changes. Compared to the nearly zero permeability value of the normal oBRB, the groups treated with ML228 and the mixture of TNF-α and TGF-β, which exhibited severe RPE morphological changes, showed significantly increased values of 2.5 x 10^-6 cm/sec and 5.8 x 10^-6 cm/sec, respectively (Fig3.C). In the case of NaIO₃, which had relatively stable ZO-1 expression, the increase was less compared to the other two treatment groups, but it was still significantly higher than normal. The TEER values of the oBRB treated with the mixture of TNF-α and TGF-β were the lowest we measured, at 22.6 Ω·cm², a reduction of about 73% compared to the control value (Fig3.D). In comparison, NaIO₃ treatment showed a 44% decrease at 46.7 Ω·cm², which was the smallest reduction we observed. These results corresponded well with the overall trends observed in the Zo-1 staining results. However, the difference in TEER values between the ML228-treated and TNF-α and TGF-β-treated groups, which had already sustained significant tight junction damage, was relatively small, suggesting the TEER loss by ML228 had already reached the maximal level, and further damage was not detectable by TEER measurement.

### 2.4 Molecular Characterization of the Damage-inducer Treated oBRB Model

The RPE cells are known to be the main source of VEGF to maintain choriocapillaris. VEGF is also known to be associated with several ocular pathologies, such as wet AMD. To determine the effects of RPE damaging agents on neovascularization, we analyzed the effects on VEGF and PEDF at the gene expression levels by qPCR and secreted protein by ELISA. The VEGF and PEDF secretion were measured using ELISA from collected cultured medium separately from the Side 1 /2 reservoirs from the RPE single model or the EC: RPE co-culture model (oBRB) that were treated with different types of damage inducers, and the ratio of VEGF and PEDF was compared to the control. The levels of VEGF and PEDF in the apical side (Side 2) were slightly higher than that of the basal side (Side 1), which is consistent with the results previously reported using ARPE-19 cells[23–26]. The gene expression and protein secretion of VEGF in the oBRB model and the RPE alone model showed increases in all the groups treated with the damage inducers, and the increase patterns for each inducer were similar (Fig 4). The most significant increase in VEGF expression was seen in the combination treatment of TGF-β and TNF-α (Fig 4A, B.). The pattern of PEDF gene expression differs between the RPE single model and the oBRB model. The results may be due to the fact that mRNA is exclusively extracted from the RPE cells in the RPE single model. But, it is extracted from both endothelial cells and RPE from the oBRB model (Fig 4A, B). As for ELISA, the combination treatment of TGF-β and TNF-α increased the PEDF level compared to the control group on the apical side (Side 2), but ML228 or NaIO3 treatment decreased PEDF compared to the control group. As a result, the VEGF/PEDF ratio increased most by ML228 and lowest by NaIO₃. On the other hand, on the basal side, the ratio of VEGF/PEDF was the highest in the combination treatment of TGF-β and TNF-α compared to the control because the secretion of PEDF was reduced in all the damage inducer treatment conditions compared to the control group (Fig.4G,H).

**Fig. 4.**
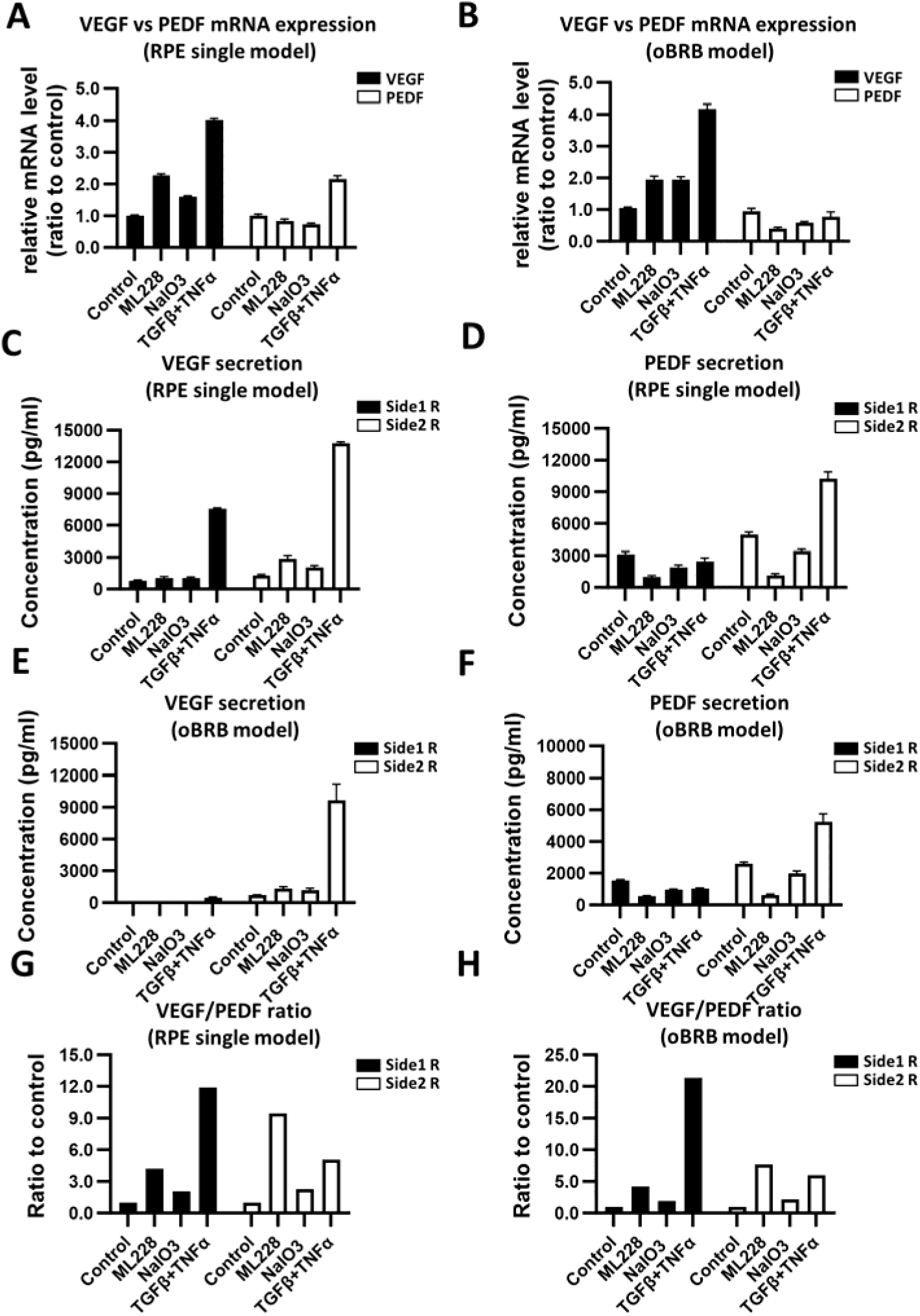
Damage inducer regulating VEGF and PEDF level in the RPE single or oBRB (REP:EC co-culture) model. (A) Normalized qPCR of VEGF and PDEF showed an increase in VEGF expression and decreased PEDF in general. The oBRB model was treated with each damaged inducer for 96 hours. Each bar represents the average of 2 replicates with standard deviation (B) Normalized qPCR results of VEGF and PEDF in the oBRB model (C) Secreted VEGF measurements using ELIA in the RPE single model (n=2) showing increase of VEGF level (D) Secreted PED measurements using ELISA in the RPE single model (n=2) showing general decrease except the apical secretion in the TGF-β and TNF-α treatment(E) Secreted VEGF measurements using ELIA in the oBRB model showing increas of the secreted level in the oBRB model (F) Secreted PEDF measurements using ELISA in the oBRB model showing general decrease by each damage inducer except the apical secretion in the TGF-β and TNF-α treatment (G) VEGF/PEDF ratios in the RPE single model damage inducer. The secretion level of VEGF was divided with PEDF and then normalized to the control condition. (n=2-3). The ratio was increased with each damage inducer treatment.

It is well known that inflammation can contribute to damage to retinal tissue and the complicated disease progression of AMD. To investigate the gene expression of IL-1β, IL-6, and IL-8 in the RPE cells, which are representative inflammatory cytokines, the gene expression by qPCR was measured in the co-culture model along with the RPE alone model. All the damage inducers increased the gene expression of IL-1β, IL-6, and IL-8 compared to the control group. The IL-1β and IL-8 expressions were increased in similar patterns in the RPE-alone model and the co-culture model compared to the control group (Fig.5A). In contrast, the expression of IL-6 was not markedly increased in the co-culture model compared to the RPE alone model. This might be explained by the role of endothelial cells in counteracting the increase of IL-6 expression by an undetermined mechanism (Fig. 5B). Interestingly, TNF-α, a regulator of inflammation, did not show much effect on the cytokine expression at a 96-hr treatment; therefore, the kinetics of the expression were investigated. The combination treatment of TGF-β and TNF-α increased IL-1β, IL-6, and IL-8 expression dramatically at 6 hours of treatment, and the expression levels decreased over time (Fig.5C, D, E). On the other hand, the ML228 and NaIO₃ treatment induced the expression of IL-1β and IL-6 lower than the combination treatment of TGF-β and TNF-α, but the expressions of these cytokines steadily increased over time. (Fig.5C, D, E)

**Fig. 5.**
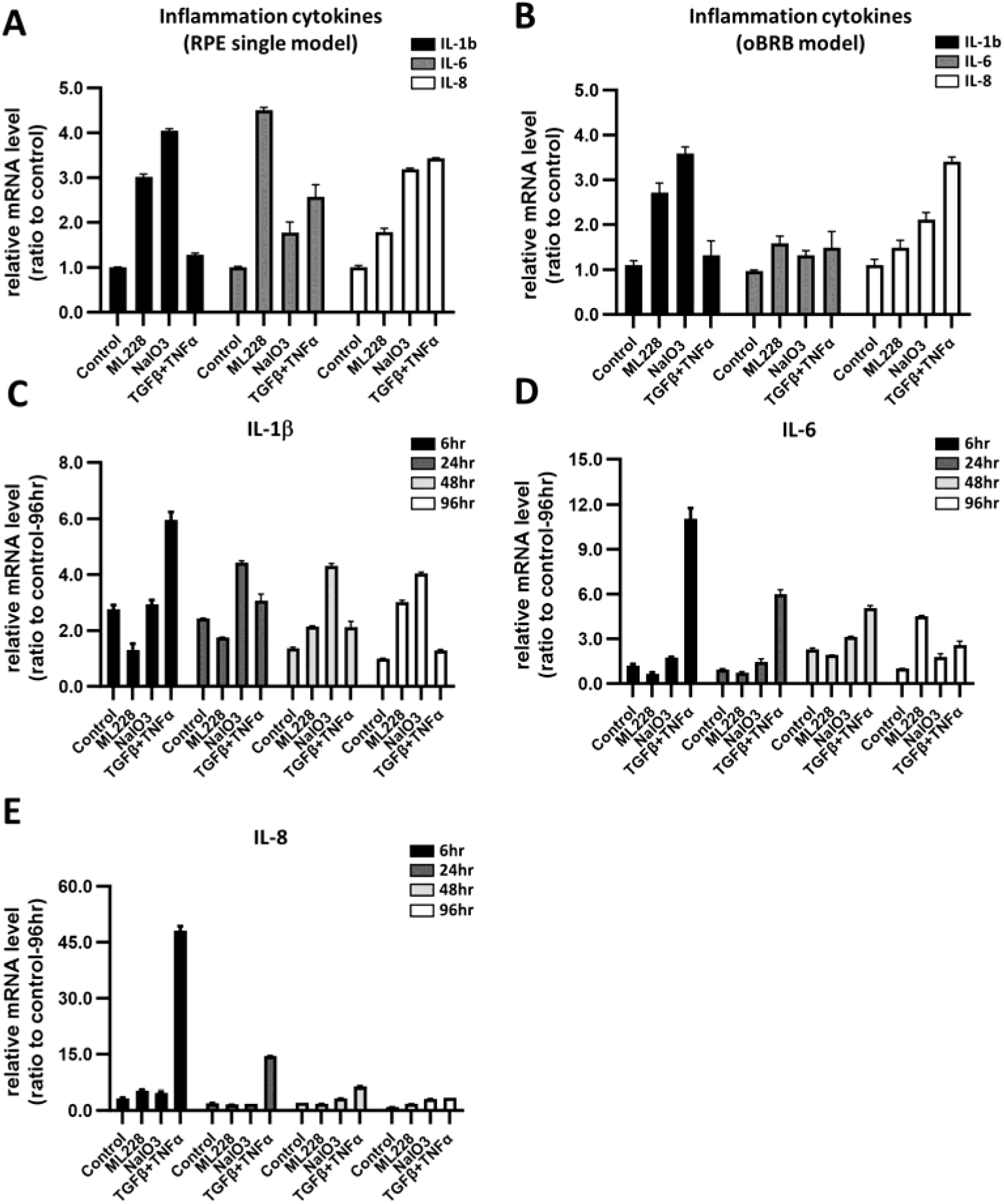
Increases of inflammation cytokine expression, IL-1b, IL-6, and IL-8 by the damage inducers (A) qPCR analysis of IL-1b, IL-6, IL-8 mRNA expression showing increases by each damage inducer post 96-hour treatment in the RPE single or (B) oBRB (EC:RPE co-culture) model. The time-dependent gene expression analysis of IL-1b(C), IL-6(D), and IL-8(E) in RPE single model at 6hr, 24hr, 48hr, and 96hr. (n=2-3).

The severe morphological changes by TGF-β and TNF-α combination treatment strongly suggest the Epithelial-Mesenchymal Transition (EMT) of in the treated RPE cells [27]. Therefore, we measured the expression of the EMT markers SNAIL, fibronectin, and Collagen A1. The EMT marker expression increased exclusively in the combination treatment of TGF-β and TNF-αbut not in the other treatment conditions. We confirmed the expression of fibronectin and α-SMA by immunostaining, showing strong expression only in the combination treatment of TGF-β and TNF-α. (Fig.6)

**Fig. 6.**
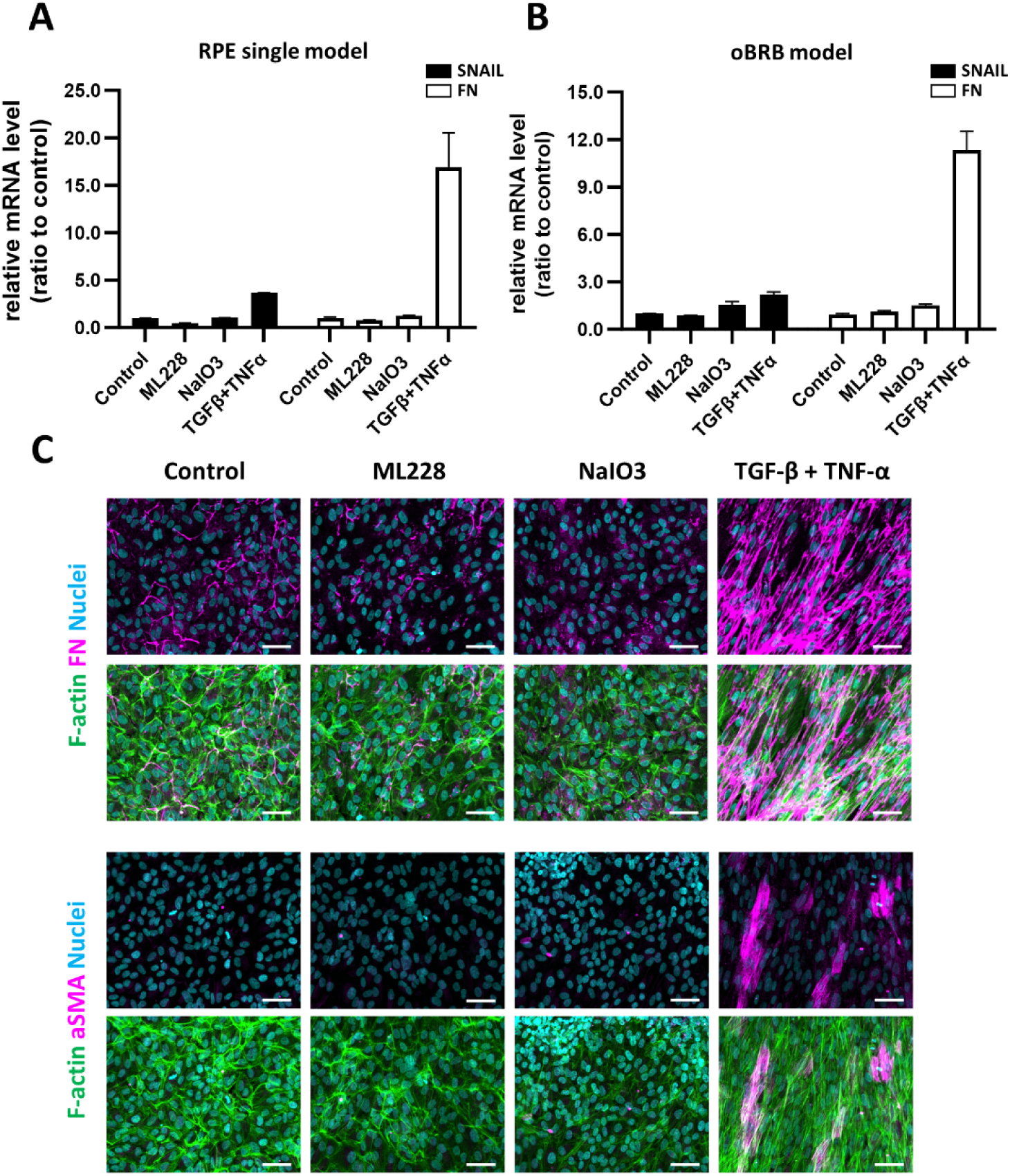
Damage inducer treatment inducing EMT marker expression in the RPE cells (A, B) qPCR analysis of Snail, Fibronectin(FN) mRNA expression in the RPE single model(A) or oBRB (EC:RPE co-culture) model (B). ARPE-19 cells in the Side2 channel were treated with ML228, NaIO₃, TGF-b, and TNF-a combination for 96hr. (C) Confocal images of F-actin and Fibronectin or a-SMA expression in damage-induced ARPE-19 cells. Both Fibronectin and a-SMA are highly expressed in TGF-b and TNF-a treated ARPE-19 cells. Scale bars: 50 μm.

To summarize, in addition to the RPE function loss, we confirmed an increase in VEGF, a decrease in PEDF, an increase in inflammation cytokines, and an increase in EMT marker expression in the damage inducer-treated oBRB model.

### 2.5 Evaluation of Anti-VEGF Antibody and Antioxidant Drug in the Damaged oBRB Model

Bevacizumab, an approved anti-VEGF drug, has demonstrated efficacy in treating wet AMD[28]. We administered this drug in our NaIO3-treated oBRB model in an attempt to reverse the observed neovascularization. We designated the group treated with NaIO₃ causing cellular oxidative stress, which prominently exhibited Choroid Neovascularization (CNV), a key characteristic of wet AMD. We treated the culture with bevacizumab only once on the fifth day of the culture, allowing full assembly of vasculature and avoiding excessive inhibition of angiogenesis by treating it too early. Bevacizumab clearly inhibited neovascularization at two different concentrations of both 5ug/ml and 50ug/ml (Fig.7.A). This effect was evident not only through the images obtained by immunostaining but also from the measured fluorescent intensity values in the middle channel, confirming the drug’s effective inhibition of CNV in our model (Fig.7.B). However, the vessel network in the Side1 channel was notably disrupted at 50 ug/ml compared to 5ug/ml. We observed bevacizumab showed a significant but partial recovery of the barrier function compared to the group treated only with NaIO₃ (Fig.7.C), but it was not extended to the full recovery of the TEER values of the control. In our model, neovascularization would not disrupt the RPE barrier since there is a hydrogel gap between the vasculature and the RPE layer; thus, we did not expect any effect on the RPE barrier function by bevacizumab. The results suggest bevacizumab may exert not only the inhibition of neovascularization but also an additional protective effect on the RPE barrier[29].

**Fig. 7.**
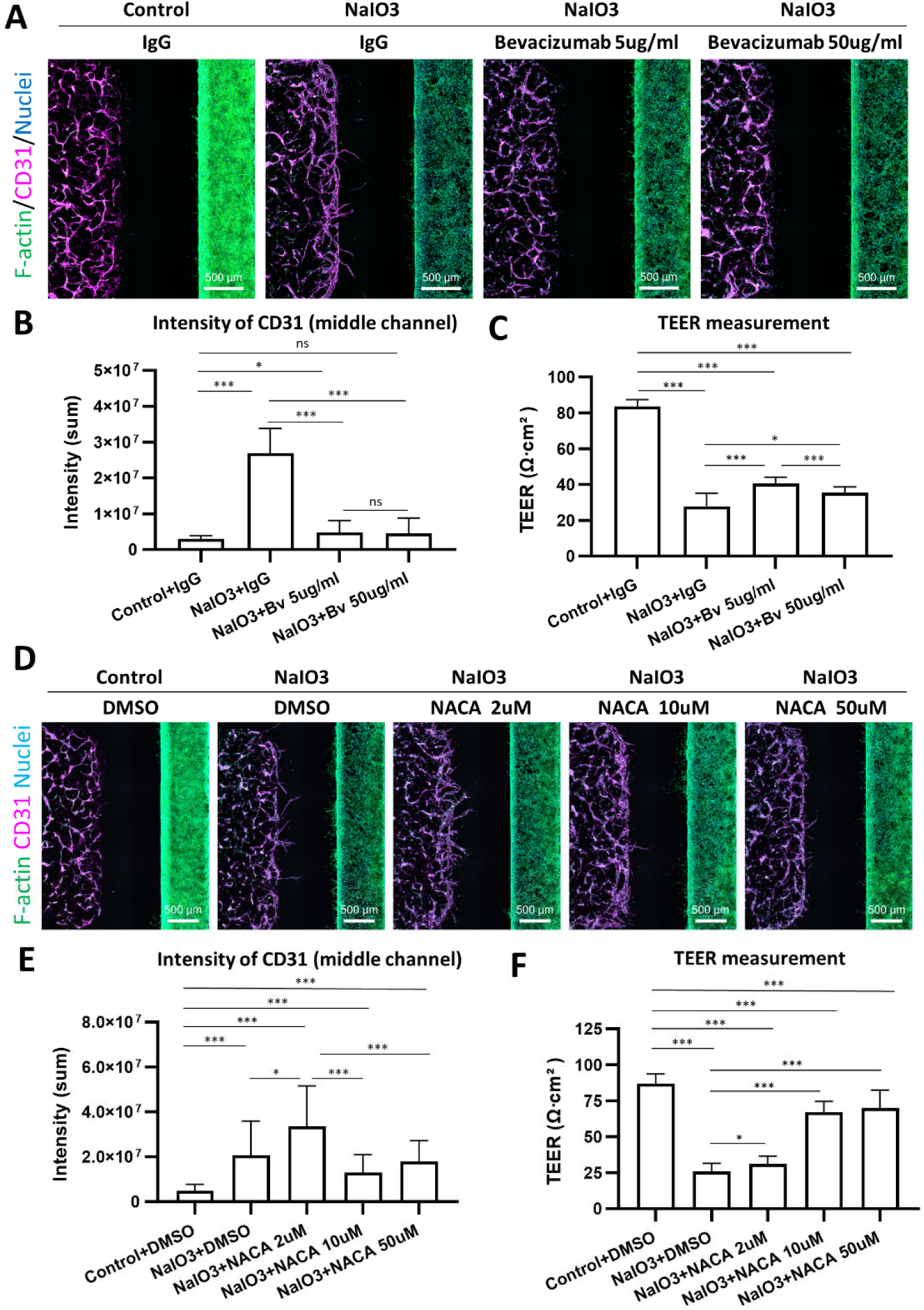
Responses of Anti-VEGF and Antioxidant drug, NACA, on NaIO3, treated oBRB model. (A) Confocal images displaying the effects of bevacizumab, anti-VEGF, treatment (0, 5, 50 µg/ml Bevacizumab or IgG) reversing new sprouting in the NaIO₃-induced oBRB model. (B) The quantification of CD31 staining intensity in the middle channel shows new sprouting by NaIO3 treatment and decreases by bevacizumab at 5 and 50 mg/ml (n=16-24). (C) TEER value decreased by NaIO3, and partially recovered bevacizumab at 5 and 50 mg/ml (n=8) (D) Confocal images depicting the response to NACA, antioxidant, treatment (0, 2, 10, 50 µM or DMSO) in the NaIO₃-damaged oBRB. (E) Quantification of CD31 staining intensity in the middle channel showing new sprouting by NaIO3 treatment and decreases by 10 and 50 mM NACA(n=15-29). (F) TEER value reduction by NaIO3 partially restored by 10 and 50 mM NACA (n=15-16)

We aimed to explore the potential of our platform for screening protective drugs. To assess their preventive efficacy in the early stages of BRB breakdown, we administered N-acetylcystein amide (NACA), an antioxidant drug with the damage inducer NaIO₃. We tested three concentrations: 2, 10, and 50 µM. While these did not reduce choroidal neovascularization (CNV) phenomena at low concentrations as dramatically as bevacizumab, we observed a decrease in CNV at the higher concentrations of 10 and 50 µM (Fig7.D). However, unlike bevacizumab, the inhibition of neovascularization by NACA was not statistically significant (Fig7.E). Conversely, at the lowest concentration of 2µM, there was a slight increase in CNV phenomena compared to the group treated with only NaIO₃. Similar trends were observed in evaluating the barrier function through TEER values. In the groups treated with NaIO₃ and 2µM of NACA, there was an approximately 75% decrease in TEER values. However, in the concentrations where CNV was reduced, the RPE recovered to about 80% of the normal oBRB control levels (Fig7.F). The effect of NACA may directly counteract the damages induced by NaIO₃ to the RPE cells and potently restore the TEER values, but the effect on inhibiting neovascularization may be achieved through protecting the RPE layer.

## 3. Discussion

Our study introduces a novel MPS specifically designed to overcome the limitations of existing approaches in modeling the outer Blood blood-retinal barrier *in vitro*. This advanced system addresses BRB pathology and evaluates RPE barrier functions while also enhancing compatibility with HTS instruments and scalability, compared to previous PDMS-based MPS[13–15]. Our efforts are currently focused on optimizing automation of the entire process to fully utilize the advantages of the 96-well plate format, which is expected to significantly improve operational efficiency and scalability, offering a stark contrast to existing models.

In our MPS, we have demonstrated stable TEER results indicative of tight junction formation in the RPE layer, even without a membrane structure or additional steps of the platform, such as tilting and rocking. It is a great advantage of avoiding the use of PDMS membranes for AMD drug discovery, which is known to absorb chemical compounds. The lateral three-compartment structure allows direct imaging of interactions between RPE and endothelial cells and sets our platform apart from traditional models that cultivate cells vertically using a membrane. Our platform supports various readout methods, such as measuring ZO-1 intensity and kinetic measurements of FITC-dextran permeability, which are critical for comprehensive barrier function assessment. This diverse array of readouts enhances the platform’s utility in high-throughput drug screening, showcasing its potential in AMD research and development.

The robust formation of the RPE monolayer, a critical component in oBRB modeling, is facilitated by our MPS’s innovative design. Key features, such as the valley-shaped Side2 channel, which supports stable barrier formation, and an open-top design for effective bubble removal, ensure barrier stability. The system’s curved structure and partially opened injection port enable precise fluid patterning and consistent cell distribution, crucial for accurately replicating the RPE layer’s dynamics and enabling detailed insights into BRB function and dysregulation that leads to AMD progression.

This MPS technological leap translates to significant biological application improvements. It enables quantifiable barrier function evaluation through TEER, ZO-1 intensity measurement, and FITC intensity measurements from FITC-dextran permeability assay and facilitates studies that require an investigation of the crosstalk between the vasculature in the choroid and the RPE barrier function, including AMD. An understanding of RPE barrier-choroidal vasculature crosstalk will open up opportunities to understand and model AMD pathogenesis. The damaging inducer treatments clearly showed the disappearance of hexagonal morphology, hallmark of intact RPE barrier, and loss of barrier function of RPE confirmed by TEER or FITC-dextran permeability. The morphological changes are tightly linked to the secretion of VEGF, which is the key mediator of the sprouting of new vessels.

Our MPS allows serial collection of media from both reservoirs to analyze secretome changes. The secretion of VEGF in polarized mature RPE is known to be higher on the basal side than the apical side, while PEDF secretion is reported to be higher on the apical side. The secretion of PEDF exhibited high levels on the apical side, consistent across both the RPE single model and the oBRB model. However, the secretion level of VEGF was higher on the apical side in both the RPE single model and the oBRB model, potentially due to the cell type ARPE-19. Previous in vitro studies have shown significantly higher levels of PEDF and VEGF secretion in polarized RPE compared to non-polarized RPE. In primary or iPSC-RPE, the secreted VEGF level is higher at the basal side, while VEGF secretion in ARPE-19 is reported higher on the apical side, which is what we observed in the oBRB model [23–26]. Moreover, notably lower levels of VEGF were observed in the oBRB model compared to the RPE single model. Results from Dr. Kapil Bharti’s lab demonstrated, via fluorescence staining, that the lower VEGF levels on the basal side in the oBRB model are due to the binding of secreted VEGF to ECs [12]. Based on these findings, the observed pattern of VEGF secretion in our oBRB model may be influenced by both the ARPE-19 cell type and the co-culture condition.

Importantly, it has been instrumental to demonstrate Bevacizumab treatment inhibited the neovascularization induced by NaIO3 suggesting new sprouting is mediated by VEGF. It still warrants investigation on how bevacizumab partially restored the barrier function where VEGF is presumably not a critical factor. Our model’s ability to provide detailed insights into the RPE barrier function and BRB dysregulation and serve as a platform for drug development underscores its potential as a transformative tool in ophthalmological research and therapy development.

However, there are limitations to consider. Firstly, the actual thickness of Bruch’s membrane, the choroid layer between the RPE and vessels, is 2-5 µm [30], whereas our platform has a distance of 1 mm, diverging from in vivo conditions. Consequently, vessel insertion to the RPE layer during neovascularization, as reported in previous literature[12, 15], could not be confirmed. There were also challenges in cultivating RPE and endothelial cells as closely as they are in vivo. Particularly, during the cultivation of the normal oBRB, endothelial migration might impede the complete formation of the RPE monolayer, which is crucial for analyzing the barrier function and neovascularization; thus, we maintained a specific middle channel distance in our chip.

Secondly, concerning the implementation of choroidal neovascularization, the well-known increase in VEGF production by the RPE, especially under pathological conditions, and its relation to neovascularization in AMD[3, 31] is critical. A decrease in PEDF expression, serving as an angiogenic inhibitor, and an increase in VEGF disrupts the VEGF/PEDF balance, promoting choroidal neovascularization[32, 33]. Under ML228 treatment, the vasculature was generally reinforced compared to the control. TNF-α and TGF-β treatment conditions led to rapid new vessel formation, but endothelial cell damage was observed later. Notably, under NaIO₃ treatment, modest vessel extension towards the RPE was observed (Fig 3). ML228 showed the highest VEGF/PEDF secretion ratio but failed to induce neovascularization in our model. Vessel sprouting towards the RPE was primarily observed in the NaIO₃ treated group, which had the least damage on the RPE barrier, the increase of VEGF expression, and the VEGF/PEDF secretion ratio. These results suggest that, for treatments other than NaIO₃, the culture period might be too short to observe neovascularization, and that neovascularization cannot be explained solely by an increase in VEGF or the VEGF/PDEF ratio, or by the concentration gradient.

Lastly, our current model using the ARPE-19 cell line is yet to fully implement a mature barrier[34]. To further develop the platform, displaying AMD phenotypes such as drusen-like characteristics and progressive phenotypes of AMD, additional optimizations are required. These include the use of more physiologically relevant cells like primary RPE cells or iPSC-derived RPE cells, optimizing culture conditions, and altering the hydrogel type to more closely replicate Bruch’s membrane.

## 4. Materials and Methods

### Fabrication of the Microfluidic Chip for Barrier Models

To enhance the adhesion of the polystyrene (PS) body fabricated via injection molding, a 10-minute plasma treatment was conducted. On the plasma-treated chip body, a double-sided pressure-sensitive adhesive tape, die-cut to match the shape of the wells, was aligned and attached. Subsequently, a polycarbonate (PC) bottom substrate was assembled. After the chip assembly, a hydrophilic surface treatment is performed using plasma treatment for approximately 20 minutes.

### Cell Culture

Human umbilical vein endothelial cells (HUVEC, Lonza) were cultured in endothelium growth medium (EGM2, Lonza), and passages 4∼6 of HUVEC were used for experiments. ARPE-19 (ATCC) cell line was cultured in DMEM/F12 (1:1) medium supplemented with 2.5mM L-Glutamine, 15mM HEPES buffer, 10% fetal bovine serum (FBS), and 1% (v/v) penicillin (100U/ml)/streptomycin (100ug/ml) (P/S), expanded to passage 30. Primary human retinal microvascular endothelial cells (HRMEC, Cell Systems) were cultured in endothelium growth medium (EGM2, Lonza), and passages 5∼7 of HUVEC were used for experiments. Human retinal pericyte cells were cultured in a pericyte medium (Sciencell), and passages 6∼8 of pericyte cells were used for experiments. All cells were maintained in a humidified incubator at 37°C and 5% CO_2_.

### Hydrogel and Cell Patterning in the Microfluidic Device

For the oBRB model, the endothelial cells in PBS were mixed with fibrinogen and aprotinin mixture solution to the concentration of HUVEC (8 million cells/ml), fibrinogen (10mg/ml) and aprotinin (0.6U/ml). 1 μl of thrombin solution (21U/ml) was dispensed into PBS. PBS (acellular) or cell suspension (8 million cells/ml) was mixed well within a tube in a ratio of 3:1 with thrombin solution. The final concentration of endothelial cells, fibrinogen, aprotinin, and thrombin was 6 million cells/ml, 2.5mg/ml, 0.15U/ml, and 1U/ml. First, patterning was started by quickly taking 1.4 μl of the acellular hydrogel solution and loading it into the specified location within the chip for middle channel patterning. After filling the middle channel patterning, it was allowed to polymerize for 6 min at room temperature. The second patterning is Side1 channel patterning with endothelial cell/fibrin mixture. 2.0 μl of endothelial cell and fibrin mixture was loaded to the Side1 channel immediately after the reaction with thrombin. For the third patterning, 10.0 μl of the medium was loaded into the Side2 channel. After complete patterning, the EGM2 was added to the Side 1 and Side 2 reservoir. The chip was incubated in a 37-degree C incubator for 24 hours. The next day, 10 μl of RPE suspension in the medium at 2 million cells/ml was loaded into the Side2 channel after removing the medium of the Side2 channel. After allowing the cells to adhere for 20∼30 minutes, 110 μl of the EGM2 to the Side1 reservoir and 175 μl of RPE differentiation medium to the Side2 reservoir were added. The RPE differentiation medium is modified from the previously established medium composition[34]. Then, the chips were incubated for 7 days to form the oBRB model, and the medium was changed every other day.

For the iBRB model, after patterning the middle and Side1 channels with acellular fibrin gel, a suspension of human retinal pericyte cells was loaded into the Side2 channel and incubated for cell attachment. Subsequently, the medium was removed. A suspension of HRMEC cells was then re-loaded into the Side2 channel. The chips were incubated in EGM2 for 7 days (Day 8), and the medium was changed every day.

### Treatment of Damage Inducers, Antioxidants, Anti-VEGF Reagents

Damage inducers, including ML228 (1uM), NaIO₃ (2.5mM), TNF-α (10 ng/ml), and TGF-β(10 ng/ml), were administered on Day 4 in the Side 2 reservoir. Damage induction with these agents was continued for 96 hours. The cells were incubated in a medium containing each damaged inducer for two days, and then the media and the damaging inducer were refreshed. After the medium was refreshed, the cells were incubated for an additional two days.

For drug testing, N-Acetyl-L-cysteinamide (NACA) was applied in the RPE medium in the Side2 reservoir with NaIO₃ (3.5 mM). For the anti-VEGF effect, a solution of bevacizumab antibody was administered to the Side2 reservoir during the second medium change in the NaIO₃-induced damage model. DMSO or IgG was used as vehicle control for NACA or bevacizumab, respectively.

### Permeability Assay

After the termination of the cell culture, the medium in both the Side1 and Side2 reservoirs was removed completely. 50 μl of RPE differentiation medium was added to Side1, and 80 μl of RPE differentiation medium with 0.1 mg/ml of 70 kDa FITC-dextran was added to the Side2 reservoir. Imaging was started immediately after the FITC-dextran solution was added. Confocal imaging was taken every 5 minutes for 30 min using a time-lapse setting. After imaging, the cells in the chip were washed with PBS and fixed with 4% paraformaldehyde solution for immunostaining. Analysis of permeability assay was done through the Open-source Microscope (OMERO) program. The channel’s Region of Interest (ROI) was selected in each well, and the intensity of FITC in ROI was measured with an analysis tool in OMERO. The co-efficiency of permeability (P_app_) is calculated with a formula.

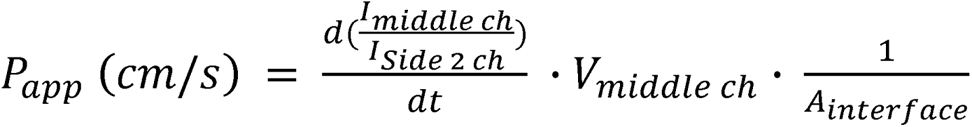

- I*_middle ch_* = Intensity of the middle channel
- I*_Side2 ch_* = Intensity of the Side2 channel
- V*_middle ch_* = Volume of middle ch (cm^3^)
- A*_interface_* = the surface area of the interface between the Side2 channel and the gel of the middle channel (cm^2^)
- t = time (sec)

### TEER Measurement

For the measurement of Transepithelial Electrical Resistance (TEER), the EVOM™ MANUAL Epithelial Volt/ohm meter (World Precision Instruments, EVM-MT-03-01) was employed. The STX4 EVOM™ ELECTRODE (World Precision Instruments, EVM-EL-03-03-01) served as the electrode for the measurements. The electrode’s longer end was positioned to reach reservoir 1, while the shorter end was set up to be situated in the Side2 channel. During this setup, the electrode on the Side1 Reservoir was positioned to be 2.4mm above the bottom. Side 2 reservoirs were replaced with an identical RPE differentiation medium, and the resistance was measured.

The resistance value, when the middle channel had an acellular gel, the Side1 channel contained endothelial cells, and the Side2 channel co-cultured with RPE, was compared to the resistance value under the same conditions but without RPE. This difference was used to compute the resistance value of the RPE. The TEER value of the RPE was determined by multiplying this with the area covered by the RPE in the Side2 Channel.

### Immunofluorescence Staining

The chip was washed 2 times with PBS, and 4% paraformaldehyde solution was added. After incubation for 20 minutes, the chip was washed 3 times with PBS. For permeabilization, 0.2% triton-X 100 in 3% BSA solution was added to both reservoirs, and the chip was incubated for 1hr ∼ 1hr 30 min at room temperature. The solution was aspirated, and a diluted antibody solution in 3% Bovine Serum Albumin (BSA) buffer was added into both reservoirs. The chip was incubated for 2 days at 4 degrees C. The chip was incubated with Alexa 647 conjugated secondary antibody for one day at 4 degrees C for the unconjugated primary antibody. For destaining, the antibody solution was removed, and the chip was washed three times with PBS and destined for more than one day. Confocal imaging was done through the CQ1 confocal device (YOKOGAWA, Japan) for anti-ZO1 (Alexa Fluor 488 or 594 conjugated, Thermos Fisher Scientific), anti-phalloidin (Alexa Fluor 488 or 594 conjugated, Thermos Fisher Scientific), and Hoechst 333342 for nuclei, anti-human CD31 and Lectin (Alexa fluor 488 or 647 conjugated, BioLegend), anti-alpha-SMA (Dako), anti-fibronectin (AbCam) and anti-PDGFRlil (Alexa fluor 488 conjugated, AbCam). For the intensity quantification for CD31 staining images for neovascularization (Fig.7A.E), an analysis script was developed and performed in OMERO. The ROI of the entire middle channel was used for the quantification of the fluorescent intensity of CD31. For each condition, we analyzed a minimum of 15 replicates, and the sum of the intensities of the samples was averaged and used for generating graphs (GraphPad Prism 10 software, San Diego, CA).

### qPCR

The medium was obliterated from the chip, and 100 μl of RLT solution (RNeasy kit, Qiagen) was added to the chip’s well. The chip was incubated for 30 min at room temperature. Cells were lysed enough using pipetting, and the lysate was transferred into the tube. Total RNA was extracted using the RNeasy kit (Qiagen) according to the instructions provided by the manufacturer. cDNA synthesis was accomplished by using M-MLV reverse transcriptase (Invitrogen). qPCR was carried out to determine the expression levels of selected genes using a StepOne plus real-time PCR system (Thermo Fisher Scientific). The primer details were listed in Table 1. The relative mRNA expression level for each gene of interest was calculated using the - 2^ΔΔCt^ method by comparing the threshold cycle value of the gene of interest to the CT value of GAPDH as a housekeeping gene.

**Table. 1.**
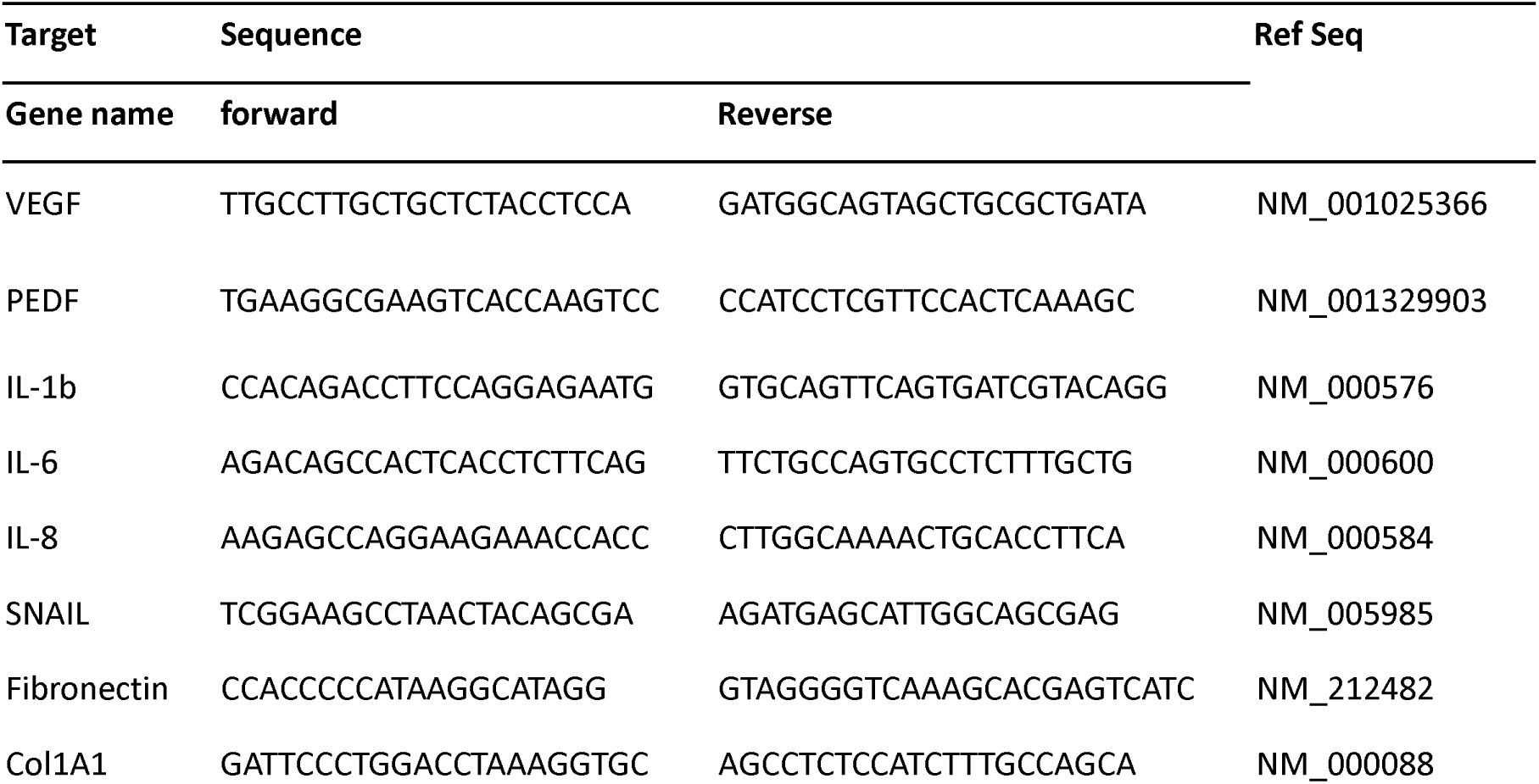
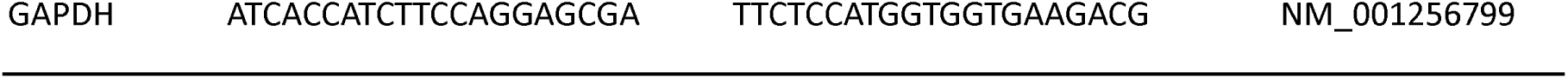
Primer sequence list for qPCR.

### ELISA

After termination of cell culture, culture medium of each reservoir was collected in a tube and stored at minus 80-degree C. VEGF or PEDF secretion level in culture medium was measured using the VEGF ELISA kit (Invitrogen or R&D systems) and PEDF ELISA kit (R&D systems) according to the instructions provided by the manufacturer. The absorbance was measured on a plate reader (Varioskan LUX, Thermo Fisher Scientific). We were using curve-fitting software to generate the standard curve. A 4-parameter algorithm provides the standard curve fit. The concentration for samples and controls is calculated from the standard curve.

### Statistical Analysis

All data of the graph were expressed as mean standard deviation. The data were analyzed with an unpaired t-test between two conditions using GraphPad Prism 10 software (San Diego, CA). At least three replicates for each condition were used for data analysis. The *p*-value of the t-test less than 0.05 was considered significant.

